# Human umbilical cord mesenchymal stromal cells ameliorate ovarian damage and restore function in Premature Ovarian Insufficiency rat model

**DOI:** 10.64898/2025.12.07.692863

**Authors:** Xue Rong, Xu Qin, Fufang Song, Yao Fu

**Affiliations:** Beauty Magic Cube Genetic and Engineering Co., Ltd., BioEngineering Research Center, Shenzhen, China; Shenzhen Spring Beauty Medical Clinic, Shenzhen, China

**Keywords:** Premature ovarian insufficiency, Umbilical cord-derived mesenchymal stromal cells, Cell-based therapy, Tissue regeneration

## Abstract

Premature ovarian insufficiency (POI) is a debilitating condition characterized by infertility and systemic health decline, with limited effective therapeutic options. This study explores the therapeutic potential of human umbilical cord-derived mesenchymal stromal cells (hUC-MSCs) in a cyclophosphamide (CTX)-induced rat model of POI. The POI rats were administered low or high doses of hUC-MSCs *via* tail vein transplantation. Ovarian function was evaluated through monitoring of the estrous cycle, serum hormone analysis, ovarian histomorphology, follicle counting, and mating assays. Immunohistochemistry was employed to assess the expression of proteins related to apoptosis and angiogenesis. Additionally, *in vitro* co-culture assays were conducted to evaluate the immunomodulatory capacity of hUC-MSCs. The results indicated that hUC-MSC transplantation restored estrous cyclicity, normalized hormone levels, increased follicle counts, and improved pregnancy outcomes in a dose-dependent manner. Histological analysis indicated a reduction in atretic follicles and an enhancement of ovarian structure. Mechanistically, hUC-MSCs were found to attenuate apoptosis by modulating the expression of Bcl-2 and Caspase-3 and to promote angiogenesis through the upregulation of VEGF. In vitro, hUC-MSCs demonstrated significant immunomodulatory effects, including the suppression of pro-inflammatory cytokine secretion and the promotion of regulatory T cell expression. In conclusion, hUC-MSCs effectively ameliorate CTX-induced ovarian damage, highlighting their promise as a regenerative therapy for POI.

## Introduction

Premature ovarian insufficiency (POI) is a clinical syndrome characterized by the cessation of normal ovarian function in women under the age of 40. The diagnosis of POI is confirmed through the presence of menstrual irregularities persisting for 4-to-6-month, alongside biochemical markers such as elevated gonadotropins and reduced estradiol levels [1–3]. A recent meta-analysis estimated the global prevalence of POI to be approximately 3.5%, with variations observed across different ethnic groups and regions, and an increasing trend over the past two decades [4]. POI significantly impacts fertility and sexual health, leading to an elevated risk of osteoporosis, cardiovascular disease, psychological issues, and increased mortality rates, thereby presenting substantial challenges for both patients and healthcare professionals [5,6]. POI is a highly heterogeneous condition with multifactorial etiologies, potentially involving genetic factors, chromosomal abnormalities, autoimmune conditions, iatrogenic factors, and environmental exposures [3,7,8]. Traditional complementary therapies, such as hormone replacement therapy, have not been effective in fully restoring ovarian endocrine and reproductive functions and may increase the incidence of other health complications [9–12]. Consequently, the development of novel and reliable therapeutic strategies to rejuvenate ovarian function is a critical objective for safeguarding and restoring reproductive health.

Advances in mesenchymal stromal cells (MSCs) research have identified MSC-based therapy as a promising approach for the treatment of POI. MSCs offer significant advantages in the field of regenerative medicine, largely attributable to their differentiation potential, paracrine activity, and immune regulatory capabilities [13–16]. These characteristics position MSCs as viable candidates for the treatment of a wide range of diseases, while simultaneously paving the way for further developments in regenerative medicine research. MSCs can be derived from a variety of sources, including bone marrow, umbilical cord, adipose tissue, dental pulp, skin, and other tissues, with biological variations influenced by their tissue of origin [17,18].

Human umbilical cord-derived mesenchymal stromal cells (hUC-MSCs) present significant advantages in the realms of cell therapy and regenerative medicine. The procurement of hUC-MSCs is both painless and non-invasive, thereby circumventing the risks and ethical concerns associated with invasive procedures typical of bone marrow-derived MSCs [19,20]. Moreover, hUC-MSCs exhibit superior proliferative capacity, minimal immunogenicity, and robust immunoregulatory properties, rendering them particularly promising for clinical applications [21,22]. Empirical studies have demonstrated that hUC-MSCs facilitate tissue repair and regeneration through the secretion of various bioactive molecules, a mechanism that is integral to wound healing and nerve injury recovery [23]. Furthermore, the incorporation of hUC-MSCs in tissue engineering has been shown to significantly enhance cell survival rates and regenerative capacity, thereby offering novel therapeutic strategies for tissue regeneration [24].

The current study on the efficacy of hUC-MSCs in enhancing ovarian function and addressing POI is presently inadequate. Although hUC-MSCs have been employed in some clinical trials for the treatment of POI [25,26], the mechanisms underlying their restorative effects remain incompletely elucidated. In this study, we established a rat model of POI induced by cyclophosphamide (CTX) injection and utilized tail vein transplantation of hUC-MSCs to validate their potential as a cell-based therapeutic approach for POI. Our findings indicate that hUC-MSC treatment restore ovarian function and reproductive capacity of rats with POI and that the therapeutic efficacy is proportional to the concentration of hUC-MSCs administered. In this study, we conducted a preliminary investigation into the potential mechanisms by which hUC-MSCs may enhance ovarian function, thereby providing a theoretical foundation for the clinical application of MSCs in the treatment of POI.

## Material and Methods

### Isolation and culture of hUC-MSCs

The umbilical cord tissue was processed to isolate the human umbilical cord mesenchymal stromal cells using a good manufacturing practice (GMP)-compliant protocol. Healthy, full-term umbilical cords were collected and immediately transferred to a sterile environment, washed, and cut into segments. The Wharton’s Jelly was extracted, washed, and centrifuged at 850 × g for 5 minutes. The gelatinous tissue was resuspended in a small volume of complete medium (MSC serum-free medium (GMP grade), Yocon, China). It was minced and placed in a T75 culture flask, then incubated at 37°C with 5% CO_2_. After 10-11 days, a substantial outgrowth of fibroblast-like cells was observed around the fragments. The medium was changed, and original fragments were removed. Once cells reached 85%-90% confluence, they were digested with gentle digestive enzyme (Yocon, China) and passaged. P4 cells were used for all subsequent studies.

### Identification of hUC-MSCs

Fourth-passage hUC-MSCs in good growth condition were harvested. The cell concentration was adjusted to 1×10^7^ cells/mL. A 100 μL aliquot of the cell suspension was transferred into individual flow cytometry tubes. For the experimental groups, the tubes were stained with fluorescent-labeled mouse anti-human monoclonal antibodies (Biolegend, USA) targeting specific markers: positive markers (CD105, CD90, CD73) and negative markers (CD45, CD34, CD19, CD11b, HLA-DR). Corresponding control groups were established using appropriately labeled mouse isotype control antibodies (Biolegend, USA). All samples were incubated at 4°C for 30 minutes, protected from light. After incubation, the cells were washed, resuspended, and then immediately analyzed using a flow cytometer (CytoFLEX S, Beckman Coulter, USA). Data analysis was performed using FlowJo software.

The cell phenotype was considered consistent with standard hMSC criteria when the following conditions were met: the positive expression rates for CD105, CD90, and CD73 were all ≥ 95.0%, while the positive expression rates for CD45, CD34, CD19, CD11b, and HLA-DR were all ≤ 2.0%.

### Identification of multi-differentiation of hUC-MSCs

hUC-MSCs in good growth condition were harvested and separately seeded into appropriate culture plates. An experimental group and a negative control group were established. The experimental group received a specific induction medium (Gibco, USA). In contrast, the control group continued to be cultured with the standard complete medium. This experimental design was implemented to validate the multi-lineage differentiation potential of the hUC-MSCs.

### Establishment of hUC-MSCs and PBMCs co-culture system

To evaluate the immunomodulatory function of UC-MSCs, the co-culture system of hUC-MSCs and PBMCs was constructed. PBMCs were first isolated from the peripheral blood of healthy volunteers and then seeded in 96-well plates at a density of 2×10^5^ cells per well. The co-culture group contained PBMCs with hUC-MSCs at a 5:1 ratio. All cells were cultured with complete RPMI 1640 medium (Thermo Fisher Scientific, USA) in incubator at 37°C with 5% CO_2_.

### Establishment of POI rat model and treatment with hUC-MSCs

All rats were obtained from Zhuhai BesTest Bio-Tech Co., Ltd (China) and underwent vaginal smearing for five consecutive days. 72 SD rats with regular estrous cycles were selected and randomly divided by body weight into a normal control group (NC group, n=16) and a model group (n=56). Rats in the model group received an intraperitoneal injection of cyclophosphamide (CTX; Medmol, China) at 10 mL/kg body weight (Fig. 1). The initial dose was 50 mg/kg. From day 2 to day 15, the dose was maintained at 5 mg/kg. Rats in the NC group received an equal volume of Saline intraperitoneally instead. All injections were administered once daily for 15 days. On day 15, 48 rats that successfully met the POI modeling criteria, based on estrous cycle disruption, were selected and randomly divided into three groups (n=16 per group): the CTX model group (CTX group), the CTX + low-dose hUC-MSCs group (1×10^6^ cells/mL, CTX + L group), and the CTX + high-dose hUC-MSCs group (3×10^6^ cells/mL, CTX + H group).

**Figure 1.**
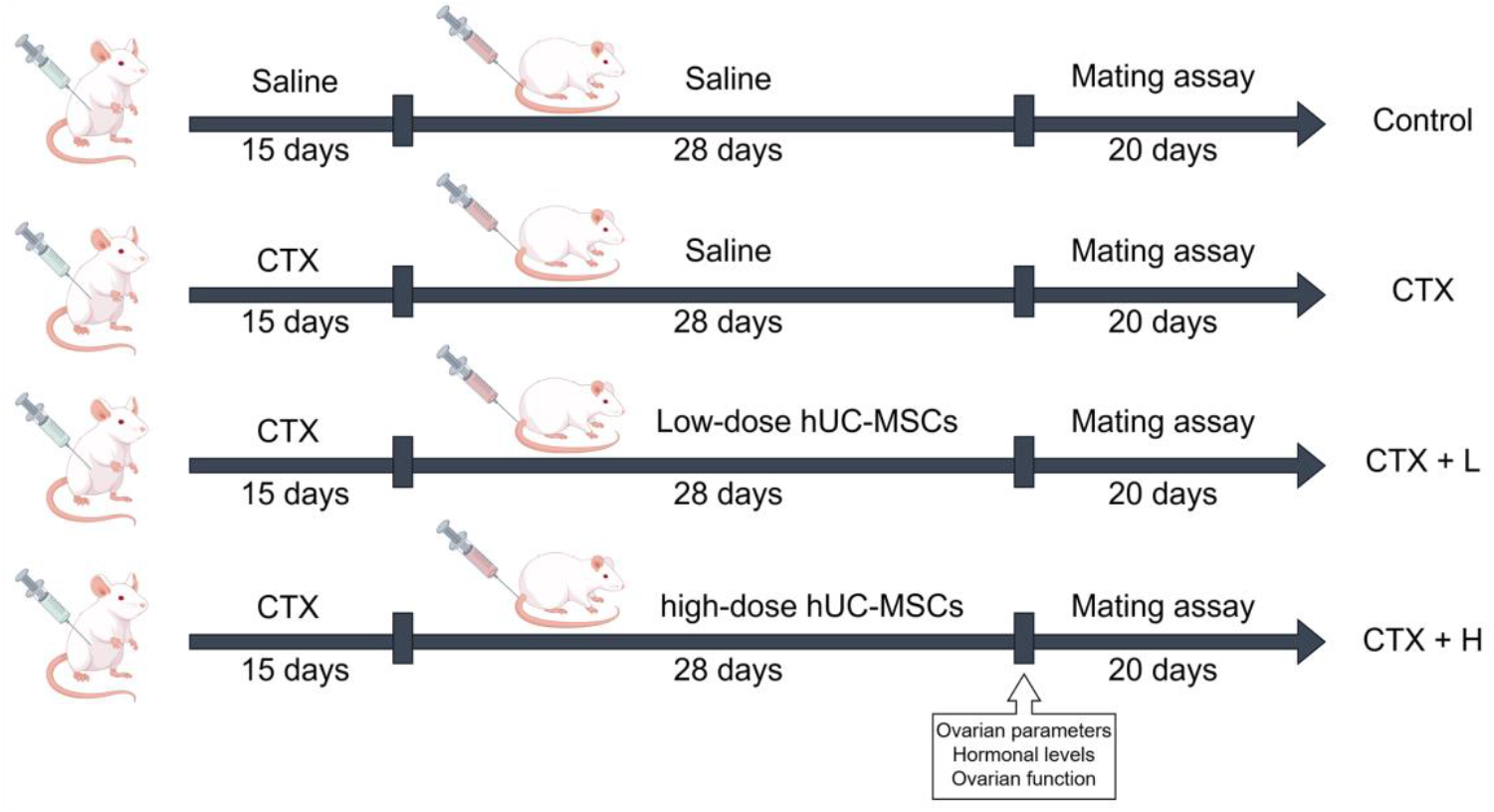
Schematic description of the experimental design. CTX (50 mg/kg for first day and 5 mg/kg/d for last days) was administered by intraperitoneal injection for 15 days. Rats in the NC group received an equal volume of normal saline intraperitoneally instead. Following CTX injection, low-dose hUC-MSCs (1.0 × 106) or high-dose hUC-MSCs (3.0 × 106) or Saline were transplanted by tail vein injection, twice a week for 4 consecutive times.

Following successful modeling and grouping, rats received their respective treatments *via* tail vein injection according to the dosing schedule. Injections were administered twice weekly for a total of eight injections. The NC group received an equal volume of Saline instead. General conditions were monitored and recorded, which include general activity, mental state, urine and feces, gait, and coat gloss. Vaginal secretions were collected daily to track changes in the estrous cycle and body weight and food intake were measured weekly.

After the final administration, ovarian tissues were harvested to perform ovarian histomorphological analysis (4 groups, n=8/group, total N=32). Both ovaries from each rat were weighed and the organ coefficient was calculated using the following formula: Ovary coefficient = [Ovarian Wet Weight (mg) / Rat Body Weight (g)] × 100%.

### ELISA analysis

Blood samples were collected from all rats at two time points: on day 15 of the modeling phase and 24 hours after the final cell administration. The samples were then promptly centrifuged with 1600 × g for 10 minutes at 4°C. The resulting plasma was carefully separated for subsequent analysis. The plasma levels of Follicle-stimulating hormone (FSH), Estradiol (E2), and Anti-Müllerian hormone (AMH) were quantitatively determined using the ELISA kits (ExCell Bio, China), following the manufacturer’s instructions.

### Hematoxylin and eosin (HE) staining

The right ovary was fixed in neutral buffered formalin and then embedded in paraffin and sectioned into 4 μm thick slices. These sections were stained with Hematoxylin and Eosin. The morphological changes in the ovarian tissue, as well as the morphology of follicles at various stages, were examined under microscope (IX 710, Olympus, Japan). A quantitative count of follicles at each developmental stage was also performed.

### Immunohistochemistry staining

The protein expression of key apoptotic and angiogenic regulatory factors (Bcl-2, Caspase-3, and VEGF) was assessed in ovarian tissue using immunohistochemistry. Paraffin sections were first deparaffinized, rehydrated, and incubated with citrate buffer for antigen retrieval. Next, the sections were incubated with a 3% hydrogen peroxide solution and blocked with 5% bovine serum albumin. Subsequently, they were then incubated with respective primary antibodies (Abcam, USA) at 4°C overnight. Following this, the sections were treated with corresponding HRP-labeled secondary antibodies (Dako, Japan) for one hour at room temperature. Color development was achieved using a diaminobenzidine (DAB) substrate kit (Abcam, USA). All images were observed and captured under microscope (IX 710, Olympus, Japan) from five non-overlapping randomly selected fields from each slide. Quantitative analysis of positive expression was performed using ImageJ software.

### Mating assay

24 hours after the completion of the female intervention therapy, eight female rats were randomly selected from each group. These females were housed with 10-week-old healthy male rats at a 1:1 ratio daily. The presence of a vaginal plug or spermatozoa in the smear was marked as a successful pregnancy. Successfully impregnated females were then housed individually and removed from the mating cycle. Twenty days after the final co-housing session, a cesarean section was performed on all females. The pregnancy status in each group was examined. The number of offspring was counted, and the embryo diameter was measured.

### Statistical analysis

All data are presented as Mean ± Standard Deviation. Statistical analysis was performed using SPSS 21.0 software. For measurement data, one-way ANOVA was applied when the variance was homogeneous. If the variance was not homogeneous, the rank sum test was used instead. For ordinal data, percentages, or ratios, the rank sum test was also employed for analysis. A *P*-value ≤ 0.05 was considered statistically significant. Other detailed information is provided in the Supplemental Information file.

## Results

### Characterization of hUC-MSCs

The umbilical cord tissue was processed in accordance with a GMP-compliant protocol to isolate and expand human umbilical cord mesenchymal stromal cells (hUC-MSCs). The cells were cultured up to the fourth passage and characterized based on their morphology, MSC surface markers, and differentiation potential. The cultured hUC-MSCs displayed a spindle-shaped, fibroblast-like morphology. Flow cytometry was utilized to confirm the phenotypic characteristics of the MSCs, which targeted specific surface markers of MSCs (CD105, CD90, CD73) and hematopoietic and endothelial markers (CD45, CD34, CD19, CD11b, HLA-DR). The analysis revealed that MSC biomarkers were strongly expressed, with an expression rate of ≥95%, while the hematopoietic and endothelial markers were minimally expressed, with an expression rate of ≤2% (Fig. 2A). These results are consistent with internationally recognized standards for MSC surface markers, thereby validating their use in subsequent studies. Furthermore, the differentiation potential of hUC-MSCs was assessed following specific *in vitro* induction protocols. The results of Oil Red O, Alizarin Red, and Alcian Blue staining confirmed the ability of hUC-MSCs to differentiate into adipocytes, osteoblasts, and chondrocytes respectively (Fig. 2B).

**Figure 2.**
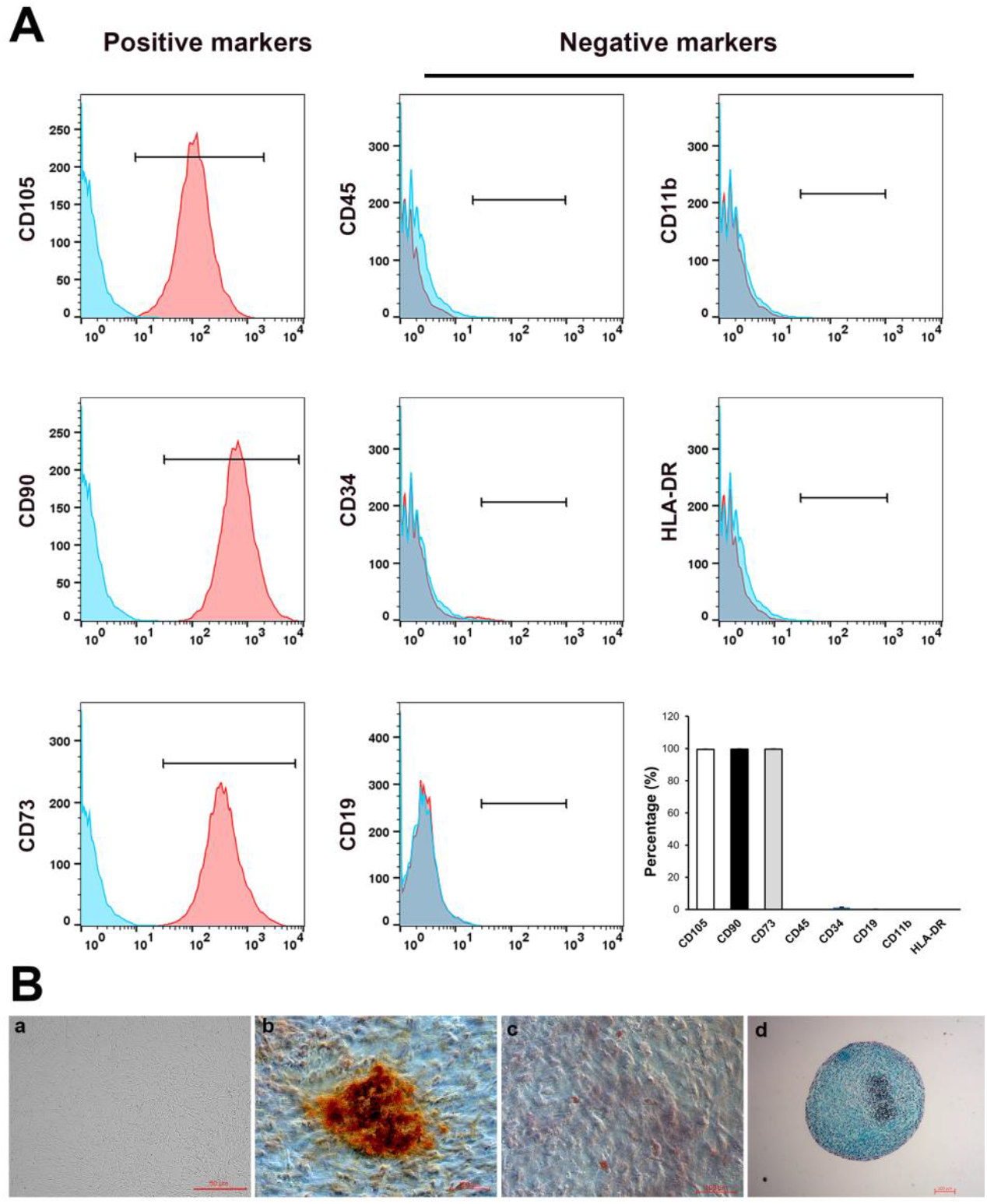
Isolation and identification of hUC-MSCs. (A) hUC-MSCs showed strong positive expression of CD105, CD90 and CD73 (>95%), and negative expression of negative cocktail including CD45, CD34, CD19, CD11b, and HLA-DR (<2%). (B) The isolated hUC-MSCs exhibited typical fibroblast-like morphology (B-a; Scale bar = 50 μm). Induction of hUC-MSCs into osteoblast (B-c; Scale bar = 100 μm), adipocytes (B-d; Scale bar = 100 μm), and chondrocytes (B-d; Scale bar = 200 μm).

### hUC-MSCs modulate immune function in vitro

To assess the immunomodulatory capabilities of hUC-MSCs, we constructed a co-culture system comprising hUC-MSCs and PBMCs *in vitro*. This experimental setup systematically investigated the impacts of hUC-MSCs on the proliferation of overall lymphocytes and specific T cell subsets (Th1, Th17, Treg), and the secretion of the inflammatory cytokine TNF-α. The results showed that, in comparison to the control groups, hUC-MSCs inhibited the proliferation of both overall lymphocytes and pro-inflammatory lymphocyte subsets, suppressed TNF-α expression, and facilitated the expansion of Treg cells, thereby demonstrating the potent immunomodulatory properties of hUC-MSCs (Fig. 3).

**Figure 3.**
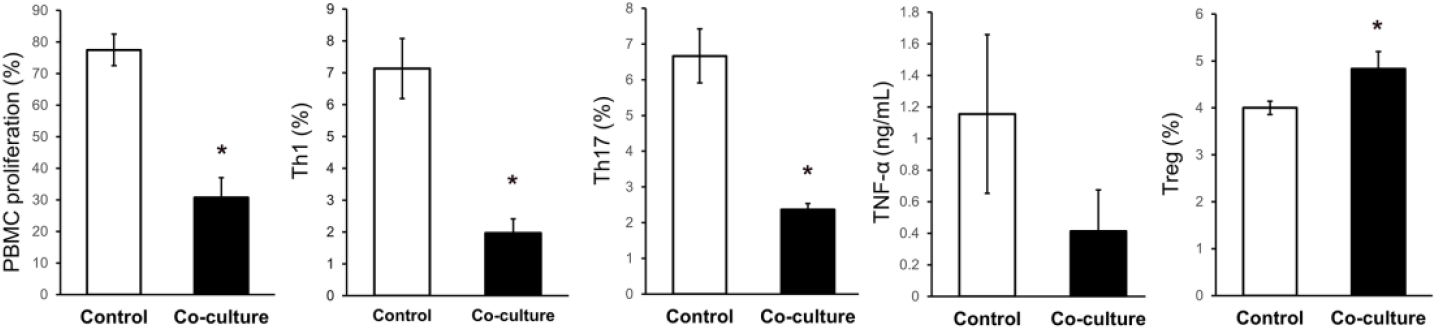
Immunomodulatory capabilities of hUC-MSCs. The impacts of hUC-MSCs on the proliferation of overall lymphocytes and specific T cell subsets (Th1, Th17, Treg), and the secretion of TNF-α were evaluated. Data represent mean ± SD, * P < 0.05.

### Establishment of CTX-induced POI rat model and transplantation with hUC-MSCs

The POI rat model was successfully established through intraperitoneal administration of cyclophosphamide (CTX) over a 15-day period (Fig. 1). Saline injection served as the normal control. Compared to the normal control group, rats in the CTX model group exhibited disrupted estrous cycles, primarily characterized by a significantly prolonged diestrus phase or persistent irregularity, ultimately leading to impaired periodic ovulation (Supplemental Fig. S1). These observations collectively confirm the successful establishment of the POI model. Following CTX administration, the rats were randomly assigned to three experimental groups, receiving either Saline or different doses of hUC-MSCs *via* tail vein injection.

Throughout the experimental period, all rats exhibited normal behavior, maintained consistent food intake, and experienced body weight gain. However, rats treated with CTX displayed significantly lower body weight and ovary coefficients compared to the normal control group. Post-transplantation, rat receiving hUC-MSCs showed mitigation of these effects, gradually approaching normal levels. Notably, high-dose hUC-MSCs treatment significantly restored both body weight and ovary coefficients compared to the CTX model group. Furthermore, the therapeutic effect observed in the high-dose UC-MSCs group was better than that of the low-dose UC-MSCs group (Fig. 4A and 4B).

**Figure 4.**
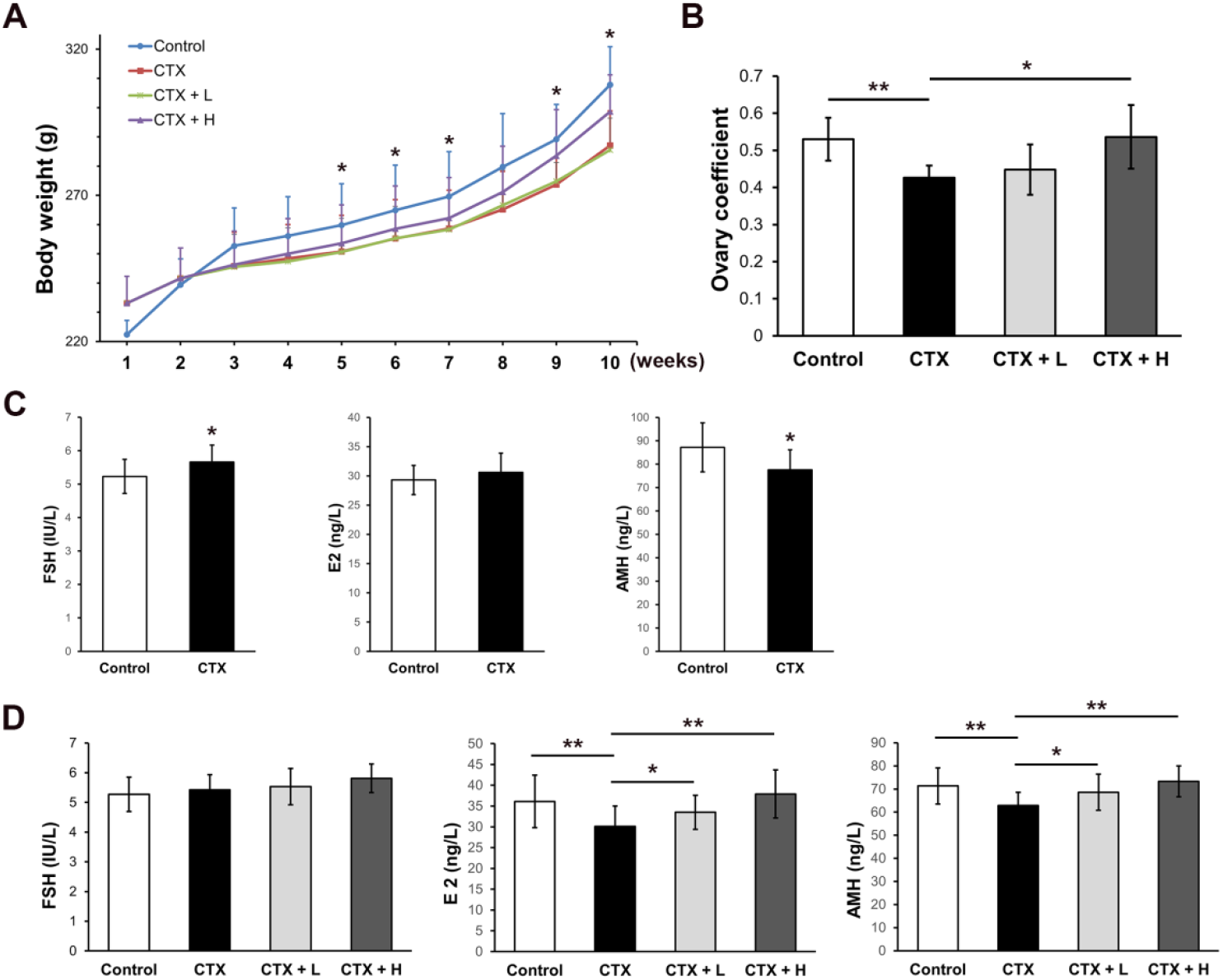
Effects of modeling and hUC-MSC transplantation on the rats with POI. (A) Changes in the body weight of rats during modeling and cell treatment were measured. * Represents P < 0.05 between the Control group and CTX group. (B) The ovary coefficient of rats in each group was determined. (C) The levels of FSH, E2, and AMH after modeling procedure. (D) The levels of FSH, E2, and AMH after hUC-MSC transplantation. Data represent mean ± SD, * P < 0.05, ** P < 0.01.

### Effect of hUC-MSC transplantation on the hormonal levels of the POI rats

To investigate the impact of hUC-MSC transplantation on ovarian function in rats with POI, analyses of serum sex hormone levels and ovarian histology were conducted. Prior to cell injection, the CTX model group exhibited significant hormonal changes. In comparison to the normal control group, the CTX model group demonstrated a marked elevation in serum FSH levels and a significant reduction in AMH levels (Fig. 4C). Hormonal profiles were reassessed 24 hours after the final cell administration. The CTX model group continued to exhibit significantly lower AMH and E2 levels compared to the normal control group. However, cell treatment effectively reversed these changes. The hUC-MSC-treated rats showed a significant increase in both AMH and E2 levels compared to the CTX model group, with the therapeutic effect being more pronounced in the high-dose hUC-MSC group (Fig. 4D).

### Restoration of ovarian morphology and function

Ovarian tissues from all experimental groups were harvested for histological examination following hUC-MSC transplantation. As depicted in Figure 5A, the ovarian morphology in the control group appeared healthy, characterized by the presence of corpora lutea and a substantial number of follicles at various developmental stages. Only a few atretic follicles were occasionally observed, and no other significant abnormalities were noted. In contrast, the ovarian histology of the CTX model group demonstrated a marked increase in the number of atretic follicles and a reduction in the number of follicles across all stages compared to the normal control group. These findings collectively confirm the successful establishment of the POI rat model and indicate that CTX injection induces damage to ovarian structure and function. Notably, following hUC-MSC transplantation, there was a restoration in the number of follicles at various stages compared to the CTX model group. Particularly in the high-dose hUC-MSC group, numerous follicles at various stages were observed, with only a small number of atretic follicles present (Fig. 5A).

**Figure 5.**
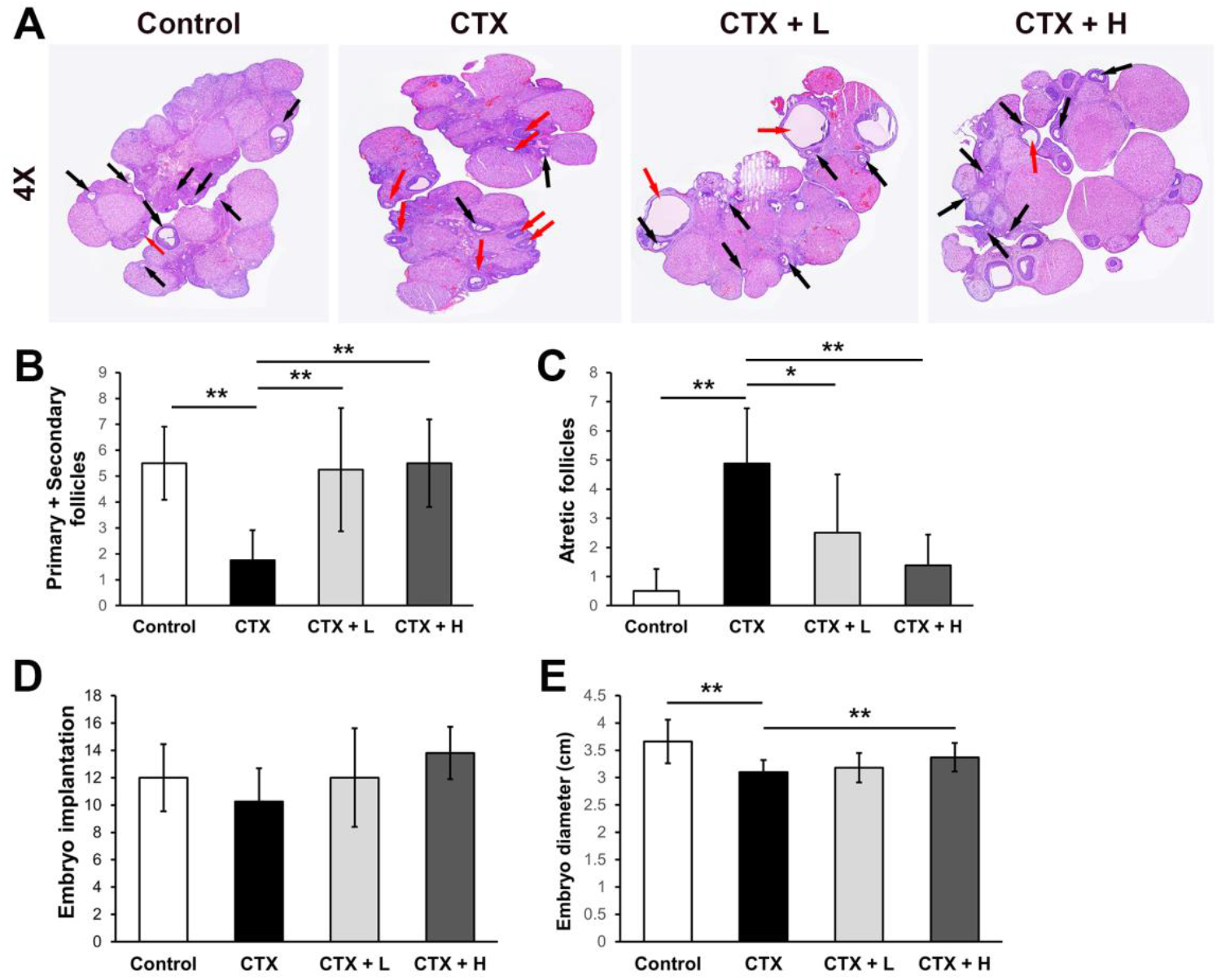
Improvement of ovarian function of rats treated with CTX by hUC-MSCs transplantation. (A) Histopathological changes of ovarian tissue by H&E staining (4 X). Black arrow represents follicles at various developmental stages. Red arrow represents atretic follicles. (B and C) The numbers of primary and secondary follicles (B) and atretic follicles (C) were statistically analyzed. (D and E) Reproductive outcomes in mating experiments. Data represent mean ± SD, * P < 0.05, ** P < 0.01.

Quantitative analysis corroborated these observations (Fig. 5B and 5C). Relative to the normal control group, the CTX model group showed a significant reduction in the number of primary and secondary follicles, alongside a marked increase in the number of atretic follicles. In contrast, both the low-dose and high-dose hUC-MSC treatment groups demonstrated significant improvements compared to the CTX model group. Specifically, there was a notable increase in the number of primary and secondary follicles and a significant decrease in the number of atretic follicles. Moreover, the improvements observed in the high-dose treatment group were substantially more pronounced than those in the low-dose group, closely approximating the levels observed in the normal control group.

These results indicate that hUC-MSC treatment effectively restored serum sex hormone levels, improved follicle counts, and reduced atretic follicles in POI rats. This suggests that hUC-MSC treatment ameliorated ovarian function and mitigated ovarian injury in rats with POI, with the effects being dose-dependent.

### hUC-MSC treatment improve reproductive capacity of rats with POI

After the final transplantation of hUC-MSCs, female rats in each experimental group were paired with fertile males. The study assessed embryo-implantation efficiency and embryo diameter to investigate the impact of hUC-MSC transplantation on the fertility of rats with POI. The CTX model group showed a reduced embryo-implantation efficiency compared to all other groups, and a significantly smaller embryo diameter relative to the normal control group. Moreover, the CTX model group demonstrated a lower pregnancy rate compared to other groups (Supplemental Table S1). Treatment with hUC-MSCs significantly ameliorated these outcomes, restoring both embryo-implantation efficiency and embryo diameter. Notably, in the high-dose treatment group, the embryo diameter was significantly greater than that observed in the CTX model group (Fig. 5D and 5E; Supplemental Fig. S2). These results indicate that hUC-MSCs can restore ovarian function and enhance the reproductive capacity of rats with POI.

### hUC-MSCs ameliorated apoptosis and promoted angiogenesis in ovarian tissue of POI rats

To elucidate the potential molecular mechanisms by which hUC-MSCs enhance ovarian function and mitigate ovarian injury in rats with POI, this study analyzed the expression of two pivotal apoptosis-related proteins, Bcl-2 and Caspase-3, alongside a critical angiogenic factor, VEGF, in ovarian tissues across all experimental groups (Fig. 6). The percentage of staining-positive areas and the mean optical density served as indicators of expression activity (Supplemental Table S2). The results indicated that, relative to the normal control group, the CTX model group showed a significant decrease in Bcl-2 expression within ovarian tissue, while Caspase-3 expression was notably elevated (Fig. 6A). Treatment with hUC-MSCs demonstrated a moderating effect. Specifically, both low-dose and high-dose hUC-MSC treatment groups exhibited a marked increase in the Bcl-2 expression compared to the CTX model group, accompanied by a reduction in Caspase-3 levels. Moreover, a significant decrease in VEGF expression was observed in the CTX model group, with both the positive area percentage and mean optical density reduced compared to the normal control group (Fig. 6B). Administration of hUC-MSCs effectively reversed this trend, as both the low-dose and high-dose stem cell groups showed increased VEGF expression compared to the CTX model group.

**Figure 6.**
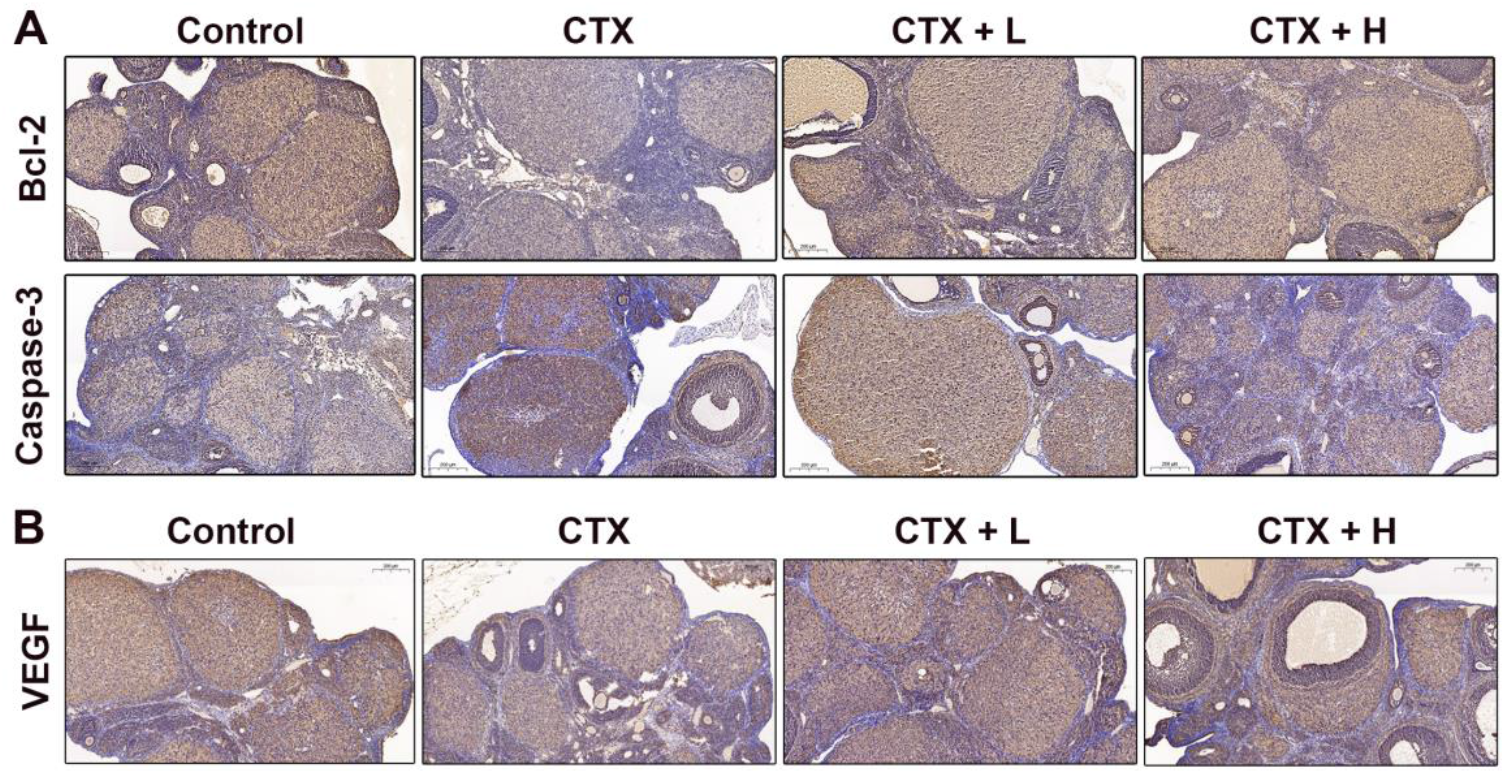
hUC-MSC transplantation ameliorated apoptosis and promoted angiogenesis in ovarian tissue of POI rats. (A) The immunohistochemistry images show the expression of apoptosis-related proteins (Bcl-2 and Caspase-3) in ovaries after hUC-MSC transplantation. (B) Representative immunohistochemistry staining images of VEGF in the ovaries of each group. Scale bar = 200 μm.

These results suggest that chemotherapy-induced POI may be associated with the decreased expression of Bcl-2 and VEGF and increased expression of Caspase-3 in the ovaries. In contrast, hUC-MSC treatment promoted Bcl-2 and VEGF expression while inhibiting Caspase-3 expression. These findings indicate that hUC-MSC transplantation has the potential to ameliorate apoptosis and promote angiogenesis in the ovaries of POI rats.

## Discussion

Our study demonstrated that the transplantation of hUC-MSCs effectively ameliorates CTX-induced ovarian damage in rats, enhances serum hormone levels, and restores ovarian function *in vivo. In vitro* assay indicated that hUC-MSCs modulate the inflammatory balance, potentially aiding in the repair of ovarian damage. Further investigations revealed that hUC-MSCs can attenuate CTX-induced cellular apoptosis, enhance ovarian angiogenesis, and promote reproductive capacity. Overall, our research elucidates the role and underlying mechanisms of hUC-MSCs in a POI rat model, providing critical insights for the development of therapeutic strategies in this field.

Umbilical cord mesenchymal stromal cells have demonstrated significant potential in the clinical treatment of various diseases through mechanisms including immune regulation, tissue repair, and regeneration [27–29]. Moreover, GMP-compliant hUC-MSCs are highly valuable for cell therapy and regenerative medicine due to their adherence to strict quality and safety standards. These standards ensure that hUC-MSCs retain their phenotypic and functional traits in animal-free media, essential for long-term culture and clinical research [30]. GMP conditions also boost their proliferation and immunosuppressive abilities, making them suitable for clinical use [31]. In clinical practice, GMP-certified hUC-MSCs demonstrate both therapeutic efficacy and proven safety [32,33]. In this work, we produced hUC-MSCs under GMP guidelines, thoroughly assessing their quality and safety (Data not shown). Meanwhile, we confirmed compliance with international cell marker standards, ensuring their therapeutic efficacy and safety [34].

In this study, the POI model was established using cyclophosphamide, a chemotherapeutic agent commonly employed in antitumor therapy [35]. Chemotherapy drugs are known to exert a range of adverse effects on the immune and reproductive systems, including the induction of granulosa cell apoptosis, vascular damage, disruption of ovarian structure, impairment of hormone secretion, and ultimately, ovarian failure [36,37]. Our results demonstrated that CTX significantly disrupted estrous cycles, reduced body weight and ovarian coefficients, and let to dysregulation of serum hormone levels (Fig. 4A and 4B). Furthermore, there was a significant decrease in the number of primary and secondary follicles, an increase in the number of atretic follicles within the ovaries of POI rats, and a consequent reduction in reproductive capacity (Fig. 5; Supplemental Fig. S2).

CTX disrupts ovarian inflammatory balance through reducing anti-inflammatory cytokines whereas increasing pro-inflammatory ones. Conversely, hUC-MSCs have been shown to positively impact tissue injury by modulating immune responses and cytokine release, thereby reducing inflammation and tissue damage [16]. In this work, we demonstrated that hUC-MSCs inhibited the proliferation of pro-inflammatory lymphocytes, suppressed the expression of TNF-α, and promoted the expansion of Treg cells, highlighting their strong immunomodulatory capacity (Fig. 3). Therefore, hUC-MSC transplantation may enhance ovarian function by creating a local anti-inflammatory microenvironment *via* secreting trophic factors that minimize further damage.

The regulation of FSH, E2, and AMH is essential for ovarian follicle development and function [38,39]. E2 is involved in the feedback regulation of FSH secretion, affecting the hormonal environment for successful reproduction [40]. AMH interacts with FSH and E2, as high FSH levels can suppress AMH expression, promoting estradiol production and follicular development [41]. In the early stages of premature ovarian insufficiency, rising FSH and E2 levels, along with decreasing AMH, indicate declining ovarian function and diminished ovarian reserve. Our data showed that body weight, estrous cycle, and hormone secretion levels in CTX-treated rats recovered to normal level after hUC-MSCs transplantation (Fig. 4). Moreover, this beneficial of hUC-MSC treatment showed dose-dependence. However, the changes in serum FSH levels after hUC-MSC treatment were less pronounced (Fig. 4D). This discrepancy could be attributed to the dynamic fluctuation of these blood-based biomarkers. Alternatively, it might result from the compensatory recovery of the rats after the initial modeling procedure. Further analysis by HE staining revealed that hUC-MSCs significantly reduced the CTX-induced ovarian follicles loss, which resulted in the restoration of reproductive capacity (Fig. 5). These results strongly support the protective effect of hUC-MSCs on the structure and function of ovarian tissue.

To further substantiate the therapeutic potential of hUC-MSCs in clinical applications, a comprehensive elucidation of their mechanisms of action is imperative. Bcl-2 and Caspase-3 are pivotal regulators of cellular apoptotic machinery, playing essential roles in maintaining cellular homeostasis and development, and are potential therapeutic targets in diseases with dysregulated apoptosis [42–44]. The interaction between Bcl-2 and Caspase-3 is also crucial for ovarian follicle development and function [45]. Bcl-2 is crucial for preventing premature apoptosis, thereby facilitating the proper development and maturation of follicles [46–48]. Moreover, Caspase-3 has been implicated in the regulation of follicular atresia and the ovarian reserve [49,50]. Our findings indicate that hUC-MSC transplantation increases Bcl-2 and decreases Caspase-3 protein expression in CTX-damaged ovarian tissue (Fig. 6A; Supplemental Table S2). The apoptosis regulation is vital for both follicle development and overall reproductive lifespan, as it affects the ovarian reserve size and fertility duration [51]. A deeper understanding of these mechanisms offers insights into fertility regulation and potential therapeutic targets for premature ovarian insufficiency and infertility.

Moreover, hUC-MSC transplantation enhances the VEGF expression in the CTX-injured sites, thereby promoting angiogenesis and tissue repair (Fig. 6B; Supplemental Table S2). This process improves the microenvironment of the injured site and lays the groundwork for addressing the incomplete treatment of POI. Studies have demonstrated that VEGF play a significant role in ovary vascularization, which is necessary for supplying nutrients and hormones to developing follicles [52–54]. Furthermore, VEGF is integral to the regulation of ovarian reserve and oocyte maturation, which are critical for successful reproduction [55]. Consequently, VEGF is a pivotal factor in ovarian physiology, influencing angiogenesis, follicular development, and overall fertility. Its expression and regulation are crucial for maintaining the delicate balance required for normal ovarian function.

This study confirmed that hUC-MSC transplantation effectively restores the ovarian function, enhances the local follicular microenvironment of ovarian tissues, and improves reproductive capability in rats with chemotherapy-induced POI. Importantly, the beneficial effects were observed in a dose-dependent manner, with high-dose administration producing results comparable to those of healthy control. Nevertheless, further comprehensive preclinical investigations are essential to validate the safety and efficacy of hUC-MSCs as a therapeutic strategy. Extensive validation clinical trials are required to establish therapeutic safety and efficacy and develop standardized treatment regimens. A thorough understanding of their efficacy across various types and severities of ovarian injury, as well as their synergistic effects with other therapeutic approaches, is necessary.

## Conclusion

Our research presents robust evidence that the transplantation of hUC-MSCs effectively reinstates ovarian function in a rat model of CTX-induced POI. The therapeutic efficacy of hUC-MSCs was evidenced by the normalization of serum hormone levels, the restoration of ovarian morphology, and the enhancement of reproductive outcomes. The recovery from CTX-induced POI is mediated by mechanisms involving anti-apoptosis and angiogenesis. These findings highlight the potential of hUC-MSCs as a promising cell-based therapeutic strategy for POI, offering a comprehensive approach to ovarian rejuvenation. Although further preclinical validation and clinical trials are necessary to establish standardized protocols and assess long-term safety, this study provides a solid foundation for the translational application of hUC-MSCs in the treatment of ovarian dysfunction and infertility.

## Ethics approval

The study was reviewed and approved by the Institutional Review Board and the Animal Care and Use Committee (Approval No. 25W166). All study participants gave written informed consent to participate in this study.

## Consent to participate

Not applicable.

## Consent for publication

Not applicable.

## Availability of data and material

Not applicable. **Code availability** Not applicable.

## Authors’ contributions

XR wrote the first draft of the manuscript; XQ and FSF contributed to review, editing, and prepare the figures; YF conceived, reviewed, and revised this paper. All authors have read and approved the final version of the manuscript.

## Acknowledgments

Figure 1 was created by Figdraw.

