## Supplement materials for "Human umbilical cord mesenchymal stromal cells ameliorate ovarian damage and restore function in Premature Ovarian Insufficiency rat model"

### Supplemental material and methods

#### Identification of multi-differentiation of hUC-MSCs

Adipogenic Differentiation: Cells were cultured in an adipogenic induction medium (Gibco, USA) for 28 days. After induction, the cells were stained with Oil Red O (Solarbio, China).The potential for adipocyte differentiation was determined by observing the formation of lipid droplets within the stained cells under a microscope.

Osteogenic Differentiation: Cells were cultured in an osteogenic induction medium (Gibco, USA) for 21 days. Following induction, Alizarin Red S staining (Solarbio, China) was performed. The potential for osteoblast differentiation was assessed by identifying the production of calcium nodules under microscopic examination.

Chondrogenic Differentiation: Cells were cultured in a chondrogenic induction medium (Gibco, USA) for 21 days. The micromasses were processed into paraffin sections and stained by Alcian blue staining solution (Solarbio, China). The potential for chondrocyte differentiation was evaluated based on the histological analysis of these sections.

#### Establishment of hUC-MSCs and PBMCs co-culture system

For lymphocyte proliferation and TNF-α secretion analysis, the setup was as follows. PBMCs were pre-labeled with 5 μM CFSE fluorescent dye (Thermo Fisher Scientific, USA). The control group (Group A) contained only CFSE-labeled PBMCs, which activated with phytohemagglutinin (PHA-M, 5 μg/mL; Thermo Fisher Scientific, USA) to induce cell proliferation. The experimental group (Group B) co-cultured PBMCs with mitomycin C-treated hUC-MSCs at a 5:1 ratio, with equal amount of PHA-M was added. A negative control group (Group C) contained only CFSE-labeled PBMCs without any stimulation. After co-culture, cells were collected to analyze the CFSE fluorescence intensity by flow cytometry (CytoFLEX S, Beckman Coulter, USA). The percentage of cell proliferation was determined based on CFSE dye dilution. Meanwhile, culture supernatants were collected from all groups and centrifuged at 3000 rpm for 10 minutes. The concentration of TNF-α was measured using a human TNF-α ELISA kit (ExCell Bio, China), following the manufacturer's instructions.

For Th1 and Th17 cell proliferation analysis, the experimental design as follows. The control group (Group A) contained only PBMCs, which stimulated with a cell activation cocktail (PIB; eBioscience, USA) for 4-6 hours at 37°C to induce intracellular cytokines. The experimental group (Group B) co-cultured PBMCs with mitomycin C-treated hUC-MSCs at a 5:1 ratio, with equal amount of PIB stimulant was added. The negative control group (Group C) contained only PBMCs without any stimulant. After co-culture, cells were first surface-stained with a CD4 fluorescent antibody (Biolegend, USA) at 4°C in the dark for 30 minutes. Intracellular staining followed. Cells were treated using a fixation/permeabilization kit (eBioscience, USA). Then, IFN-γ and IL-17A fluorescent antibodies (Biolegend, USA) were added and incubated at 4°C in the dark for 30 minutes. The proportions of Th1 ($CD4^{+}IFN\text{-}\gamma^{+}$) and Th17 ($CD4^{+}IL\text{-}17A^{+}$) cells were detected by flow cytometry.

For Treg cell proliferation analysis, the experimental group (Group A) co-cultured PBMCs with mitomycin C-treated hUC-MSCs at a 5:1 ratio. The negative control group (Group B) contained only PBMCs. After co-culture, Treg cell staining was performed. Cells were first surface-stained with CD4 and CD25 fluorescent antibodies (Biolegend, USA) at 4°C in the dark for 30 minutes. Subsequently, cells were fixed and permeabilized using the Foxp3/Transcription Factor Staining Buffer Set (Biolegend, USA). Finally, a Foxp3 fluorescent antibody (Biolegend, USA) was added and incubated at 4°C in the dark for 30 minutes. The percentage of Treg ($CD4^{+}CD25^{+}Foxp3^{+}$) cells was detected by flow cytometry.

**Supplemental Figures**

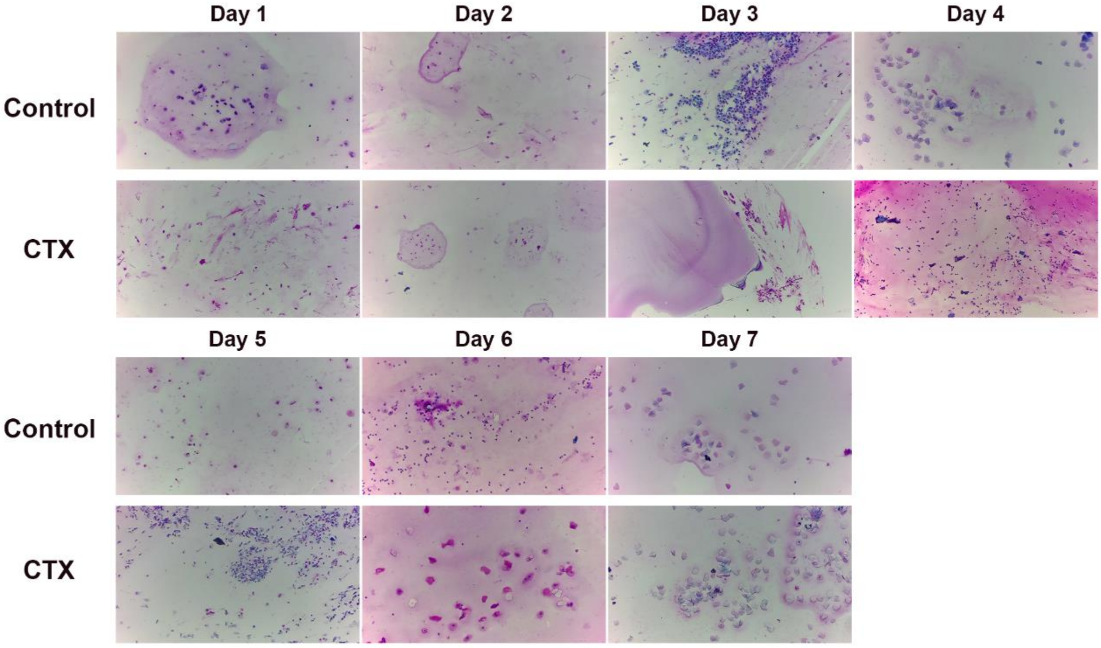


Supplemental Figure S1. Comparison of estrous cycles in the control group and CTX model group for 7 days during the modeling procedure. Compared to the normal control group, rats in the CTX model group exhibited disrupted estrous cycles, primarily characterized by a significantly prolonged diestrus phase or persistent irregularity

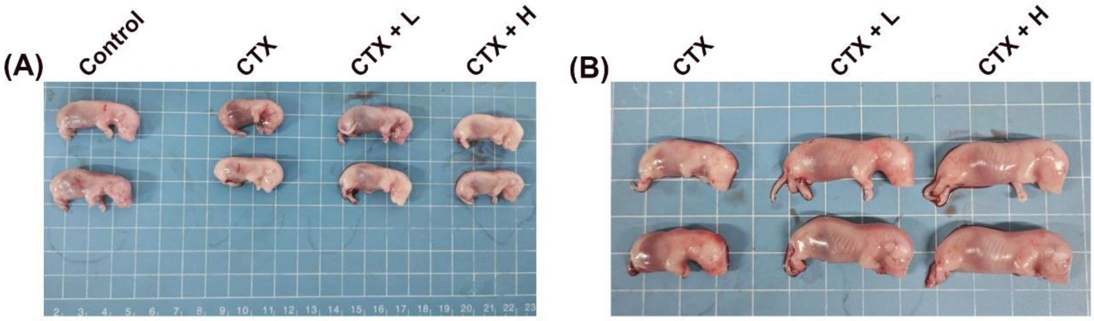


Supplemental Figure S2. Comparison of embryo diameter in different groups. (A) Comparison of embryo diameter in the control group, CTX model group, CTX + low-dose hUC-MSCs group, and CTX + high-dose hUC-MSCs group. (B) Clear comparison of embryo diameter in the CTX model group, CTX + low-dose hUC-MSCs group, and CTX + high-dose hUC-MSCs group

**Supplemental Tables**

Supplemental Table S1. The number of pregnant rats and embryo implantation and embryo diameter of the rats from all groups

| Group | Pregnancy number | Embryo implantation | Embryo diameter |
|---|---|---|---|
| Control | 5 | 12.0 ± 2.45 | 3.66 ± 0.4 |
| CTX | 3 | 10.25 ± 2.44 | 3.1 ± 0.22▲▲ |
| CTX + L | 4 | 12.0 ± 3.61 | 3.18 ± 0.27 |
| CTX + H | 5 | 13.8 ± 1.92 | 3.37 ± 0.26** |

▲▲represent P<0.01 compared to the Control group; **represent P<0.01 compared to the CTX group.

Supplemental Table S2. Expression of Bcl-2, Caspase-3, and VEGF in the ovaries from all groups

| Groups | n | Bcl-2 expression | | Caspase-3expression | | VEGF expression | |
|---|---|---|---|---|---|---|---|
| | | Positive area percentage (%) | Mean optical density | Positive area percentage (%) | Mean optical density | Positive area percentage (%) | Mean optical density |
| Control | 8 | 23.30±3.11 | 0.329±0.033 | 19.31±3.71 | 0.299±0.039 | 24.06±3.35 | 0.396±0.047 |
| CTX | 8 | 17.85±3.02▲▲ | 0.285±0.033▲ | 23.41±3.73▲ | 0.319±0.079 | 17.57±3.92▲▲ | 0.362±0.032 |
| CTX + L | 8 | 21.61±4.88* | 0.310±0.041 | 20.96±2.75 | 0.307±0.046 | 21.83±4.80* | 0.384±0.035 |
| CTX + H | 8 | 23.19±1.89** | 0.317±0.031 | 20.85±2.53 | 0.300±0.045 | 23.48±3.01** | 0.391±0.044 |

▲represent P<0.05 and ▲▲represent P<0.01 compared to the Control group; * represent P<0.05 and **represent P<0.01 compared to the CTX group.
